# SARS-CoV-2 501Y.V2 (B.1.351) elicits cross-reactive neutralizing antibodies

**DOI:** 10.1101/2021.03.06.434193

**Authors:** Thandeka Moyo-Gwete, Mashudu Madzivhandila, Zanele Makhado, Frances Ayres, Donald Mhlanga, Brent Oosthuysen, Bronwen E. Lambson, Prudence Kgagudi, Houriiyah Tegally, Arash Iranzadeh, Deelan Doolabh, Lynn Tyers, Lionel R. Chinhoyi, Mathilda Mennen, Sango Skelem, Gert Marais, Constantinos Kurt Wibmer, Jinal N Bhiman, Veronica Ueckermann, Theresa Rossouw, Michael Boswell, Tulio de Oliveira, Carolyn Williamson, Wendy A Burgers, Ntobeko Ntusi, Lynn Morris, Penny L Moore

**Affiliations:** National Institute for Communicable Diseases of the National Health Laboratory Service, Johannesburg, South Africa; Antibody Immunity Research Unit, School of Pathology, Faculty of Health Sciences, University of the Witwatersrand, Johannesburg, South Africa; KwaZulu-Natal Research Innovation and Sequencing Platform (KRISP), Department of Laboratory Medicine & Medical Sciences, University of KwaZulu-Natal, Durban, South Africa; Institute of Infectious Disease and Molecular Medicine, Division of Medical Virology, Department of Pathology, University of Cape Town, Cape Town, South Africa; Wellcome Centre for Infectious Diseases Research in Africa, University of Cape Town, Cape Town, South Africa; Division of Cardiology, Department of Medicine, University of Cape Town and Groote Schuur Hospital, Cape Town, South Africa; Hatter Institute for Cardiovascular Research in Africa, Faculty of Health Sciences, University of Cape Town, Cape Town, South Africa; National Health Laboratory Services, Groote Schuur Hospital, Cape Town, South Africa; Department of Virology, School of Pathology, Faculty of Health Sciences, University of the Witwatersrand, Johannesburg, South Africa; Division for Infectious Diseases, Department of Internal Medicine, Steve Biko Academic Hospital and University of Pretoria, Pretoria, South Africa; Department of Immunology, Faculty of Health Sciences, University of Pretoria, Pretoria, South Africa; Centre for the AIDS Programme of Research in South Africa (CAPRISA), Durban, South Africa; Department of Global Health, University of Washington, Seattle, USA

## Abstract

Neutralization escape by SARS-CoV-2 variants, as has been observed in the 501Y.V2 (B.1.351) variant, has impacted the efficacy of first generation COVID-19 vaccines. Here, the antibody response to the 501Y.V2 variant was examined in a cohort of patients hospitalized with COVID-19 in early 2021 - when over 90% of infections in South Africa were attributed to 501Y.V2. Robust binding and neutralizing antibody titers to the 501Y.V2 variant were detected and these binding antibodies showed high levels of cross-reactivity for the original variant, from the first wave. In contrast to an earlier study where sera from individuals infected with the original variant showed dramatically reduced potency against 501Y.V2, sera from 501Y.V2-infected patients maintained good cross-reactivity against viruses from the first wave. Furthermore, sera from 501Y.V2-infected patients also neutralized the 501Y.V3 (P.1) variant first described in Brazil, and now circulating globally. Collectively these data suggest that the antibody response in patients infected with 501Y.V2 has a broad specificity and that vaccines designed with the 501Y.V2 sequence may elicit more cross-reactive responses.

## Introduction

The global pandemic of severe acute respiratory syndrome coronavirus 2 (SARS-CoV-2), the virus causing coronavirus disease 2019 (COVID-19) (Zhou et al., 2020), has resulted in a concerted and highly successful vaccine development program (Krammer, 2020, Rawat et al., 2020). Almost all SARS-CoV-2 vaccines include the spike protein to elicit neutralizing antibodies (nAbs), likely the primary correlate of protection from infection (Rogers et al., 2020, Yu et al., 2020, Jackson et al., 2020, Zhang et al., 2020b, Mercado et al., 2020). Early evidence for the emergence of spike variants was seen in the rapid global selection of spikes bearing the D614G substitution, which resulted in improved binding to the host receptor, angiotensin converting enzyme (ACE-2), but did not reduce sensitivity to nAbs (Korber et al., 2020a, Korber et al., 2020b, Zhang et al., 2020a). However, more recently, three variants of concern have emerged around the world, all of which exhibit immune escape.

The 501Y.V1 (B.1.1.7) lineage, first detected in the United Kingdom, has enhanced transmissibility and an N501Y change that increased affinity for ACE-2 (Rambaut et al., 2020b, Liu et al., 2021). The 501Y.V1 variant did not show substantially increased resistance to neutralizing antibodies, though the subsequent emergence within this lineage of the E484K mutation suggests immune escape (Muik et al., 2021, Wu et al., 2021, Li et al., 2021, Graham et al., 2021, Wise, 2021). This is in contrast to the 501Y.V2 lineage (B.1.351), which was identified in South Africa in October 2020 and subsequently became the dominant circulating variant (Tegally et al., 2021). This lineage includes three changes in the receptor binding domain (RBD) (K417N, E484K and N501Y), five in the N-terminal domain (NTD) (L18F, D80A, D215G, R246I and a deletion at 242-244) and one in the S2 subunit (A701V). These confer partial to complete resistance to convalescent plasma and to some monoclonal antibodies (mAbs), including class I/II RBD-directed mAbs and 4A8 which targets the NTD (Wibmer et al., 2021, Cele et al., 2021, Wang et al., 2021b). A third variant of concern, the 501Y.V3 (P.1) lineage identified in Brazil contains similar RBD mutations (K417T, E484K and N501Y) as well as eight other changes across the spike (L18F, T20N, P26S, D138Y, R190S, H655Y, T1027I and V1176F), and also exhibits increased neutralization resistance (Faria et al., 2021, Wang et al., 2021c, Garcia-Beltran et al., 2021).

As an increasing number of variants are being detected globally, these findings have raised widespread concerns about their impact on the efficacy of current vaccines. Indeed, vaccine efficacy against 501Y.V2 was reduced in the ChAdOx1 nCoV-19 (AZD1222) trial which failed to protect against mild/moderate COVID-19 disease in people infected with 501Y.V2 (Madhi et al., 2021). Moreover, sera from Pfizer-BioNTech and Moderna mRNA vaccinees showed substantially reduced neutralization of 501Y.V2 (Wang et al., 2021b). In response, vaccine developers have moved towards replacement of the original vaccine sequence with 501Y.V2 sequences (Moderna, 2021), on the assumption that these vaccines will elicit equivalently good neutralizing responses. However, it is unknown whether the 501Y.V2 variant elicits a potent neutralizing response during infection. Similarly, the extent to which antibodies elicited by 501Y.V2 infection cross-react with other variants is not known, but has implications for the ability of second-generation vaccines to protect against infection by the original and emerging SARS-CoV-2 lineages.

Here, we characterize a large cohort of patients hospitalized after the emergence and dominance of the 501Y.V2 variant in Cape Town, South Africa, during the second wave. We show that 501Y.V2 infection elicits potent binding and neutralizing responses that exhibit cross-reactivity against viruses circulating before the emergence of 501Y.V2 variant (D614G, referred to here as the original variant), as well as the 501Y.V3 (P.1) variant first described in Brazil. Therefore, we hypothesize that an immunogen based on 501Y.V2 may elicit antibodies that can protect against multiple circulating SARS-CoV-2 lineages.

## Results

### The 501Y.V2 variant elicits potent binding and neutralizing antibodies

Blood samples from 89 COVID-19 patients admitted to the Groote Schuur Hospital (GSH) were collected between 31 December 2020 and 15 January 2021 in Cape Town, South Africa. Sequencing was performed for 28/89 (31%) patients, and all were shown to be 501Y.V2 by mutational and phylogenetic analysis (**Figure S1A**). In addition, the local epidemic in Cape Town and South Africa as a whole was dominated by 501Y.V2, which was responsible for over 90% of infections in December 2020 and January 2021 (**Figure S1B**) (Tegally et al., 2021). Furthermore, no patient reported prior SARS-CoV-2 infection.

We first assessed the magnitude of the binding and neutralizing antibody responses in the GSH patients to the 501Y.V2 spike protein (matching the presumed infecting virus). The median time post-SARS-CoV-2 PCR-positive test for the GSH cohort was 7 days (Interquartile range [IQR]: 3-11). These were compared with samples from a similar hospitalized cohort (n=62) recruited in Pretoria, South Africa in May – September 2020 during wave 1, prior to the emergence of 501Y.V2. Samples from this hospitalized cohort who were infected with the original variant, were taken at admission and a follow-up visit 5-9 days post-admission. Admission occurred at a median of 3 days (IQR:2-5) after testing, and follow-up samples were taken a median of 10 days (IQR:9-13) after testing. We observed no significant difference in the magnitude of binding (**Figure 1A**) or neutralizing (**Figure 1B**) responses between the GSH cohort and the wave 1 admission samples, but titers at the follow-up timepoint were significantly higher in the wave 1 cohort than in the GSH cohort **(Figure 1A and B)**. These differences may be due to the varying time of sampling, and the rapidity with which neutralizing antibodies mature between 3 and 10 days after testing. Thus, like the original variant, 501Y.V2 elicits high titer binding and neutralizing antibody responses.

**Figure 1.**
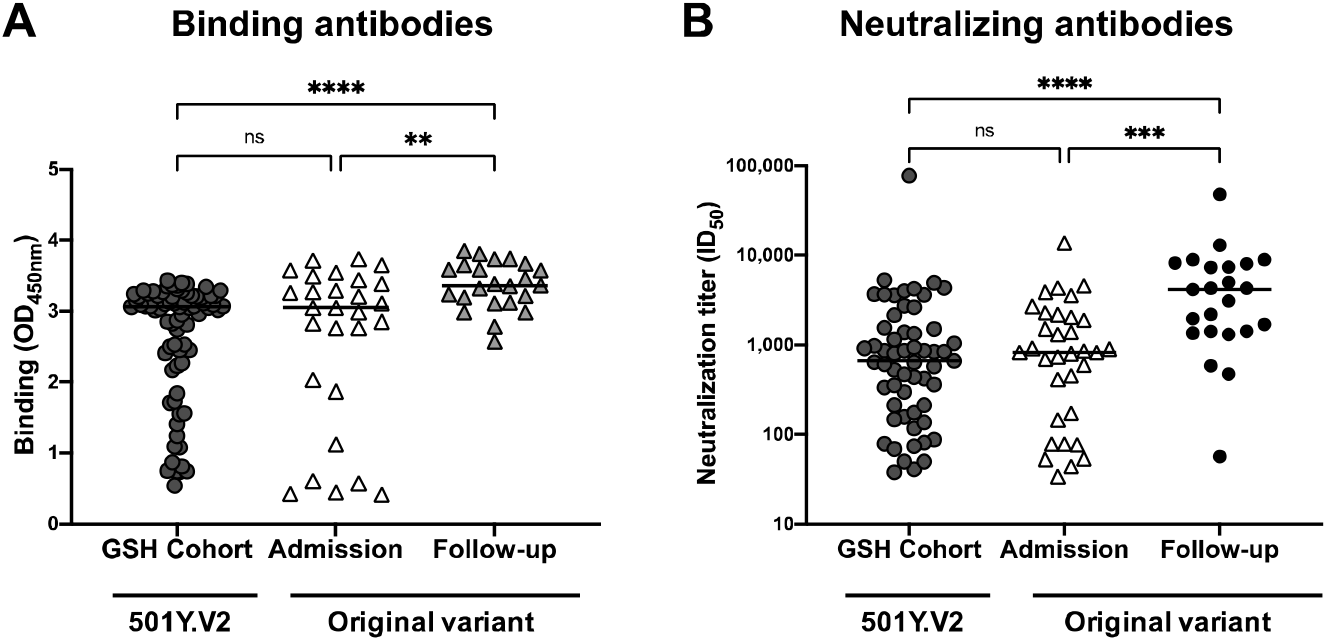
501Y.V2 elicits robust binding and neutralizing antibody responses. Plasma samples from hospitalized individuals infected with either the 501Y.V2 variant (n=89) or the original variant (n=62) were tested for (**A**) Binding (OD_450nm_) to the sequence-matched SARS-CoV-2 spike antigen and (**B**) neutralizing activity (ID_50_) against SARS-CoV-2 pseudoviruses. Binding was assessed at a single dilution of 1:100, and neutralization titers from a starting dilution of 1:20. For both binding and neutralization, only individuals with binding or neutralizing responses are shown. For the 501Y.V2 cohort, binding data are shown for 75 patients, and neutralization data for 57. For the original variant cohort, 28 samples were measured at admission and 23 at follow-up for binding responses; 33 at admission and 23 at follow-up for neutralization assays. Samples were scored as positive when binding was greater than an OD_450nm_>0.4 and the threshold of detection for the neutralization assay is ID_50_>20. All experiments were performed in duplicate. Significance is shown as: ns= p>0.05, **p<0.01, ***p<0.001, ****p<0.0001.

### Infection with 501Y.V2 elicits cross-reactive binding antibodies to RBD and full spike protein

We assessed the magnitude of binding responses to the SBD (RBD + subdomain 1) and spike protein of the original and 501Y.V2 variants. We tested 89 samples at a single dilution (1:100), and observed a strong correlation in binding to the original variant and 501Y.V2 for both the SBD (**Figure 2A**) and the full spike protein (**Figure 2B**). Titration of a subset of 46 samples revealed that although plasma samples had higher titers to the spike of 501Y.V2 than to that of the original variant (average reduction of 1.7-fold), high level binding to the original variant remained (**Figure 2C**). Thus, binding antibodies elicited by 501Y.V2 are able to cross-react with the original circulating variant.

**Figure 2:**
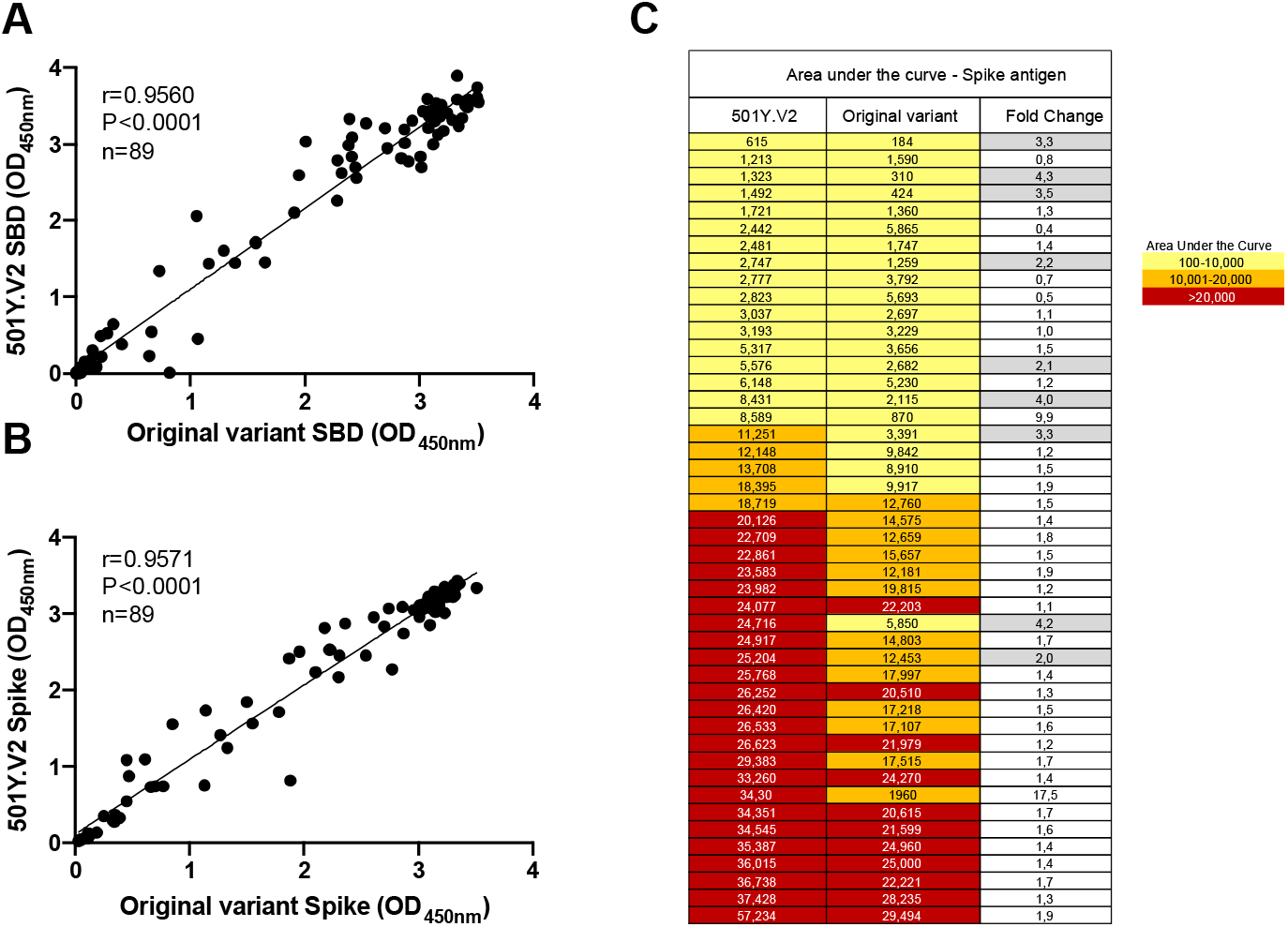
Plasma binding antibodies in 501Y.V2 infected individuals are cross-reactive. Plasma samples from 89 GSH patients were tested for binding to (**A**) SBD (original variant and 501Y.V2) and (**B**) full spike (original variant and 501Y.V2). Binding in A and B was assessed at a single plasma dilution of 1:100. (**C**) Plasma samples from a subset of n=46 individuals were titrated out in full against the 501Y.V2 spike (column 1) and the spike from the original variant (column 2). Data are plotted as area under the curve, ranked by titers against 501Y.V2, and colored according to magnitude. The fold differences between binding to each of the two lineages is shown in column 3, with grey shading denoting fold changes ≥ 2. All experiments were performed in duplicate.

### Neutralizing antibodies elicited by the 501Y.V2 variant are cross-reactive

We previously reported substantially lower neutralization of the 501Y.V2 by plasma from individuals infected with the original variant, with 48% of sera unable to neutralize 501Y.V2 (**Figure 3A, Figure S2A**) (Wibmer et al., 2021). Here we performed the reverse experiment, by assessing the cross-reactivity of neutralizing responses in the GSH 501Y.V2 cohort, against the original variant and 501Y.V3 (P.1). We first tested 57 GSH donor sera against both 501Y.V2 and the original variant (**Figure 3B**). Strikingly, 53/57 plasma samples maintained neutralization activity against the original variant, with an average fold reduction of 3 (**Figure 3B**) and a geometric mean titer (GMT) of 203 (**Figure 4A**). Two plasma samples showed the opposite trend, with higher neutralization of the original variant (>5 fold-higher), compared to 501Y.V2 (**Figure 3B**). We do not have sequencing data for these samples but it is likely that these individuals were infected with the original variant and not 501Y.V2. A construct that contains only the three RBD changes in 501Y.V2 (K417N, E484K, N501Y) showed similar titers to the full 501Y.V2 (GMT:669 compared to 501Y.V2: GMT:686, for this subset of n=51) (**Figure S2**). This suggests that much of the neutralization activity against 501Y.V2 is directed against the RBD, which is shared between these two constructs. When we limited the analysis to 22/28 donors for whom sequencing data proved infection with 501Y.V2 and who had antibody binding titers, we observed the same pattern (**Figure 3C**). Lastly, we tested a subset of 10 samples against 501Y.V3 (P.1), and also showed high levels of neutralization of this variant, with some samples showing increased potency, a finding that may be due to the very different NTD regions of these variants (**Figure 3D**) (Wang et al., 2021c).

**Figure 3.**
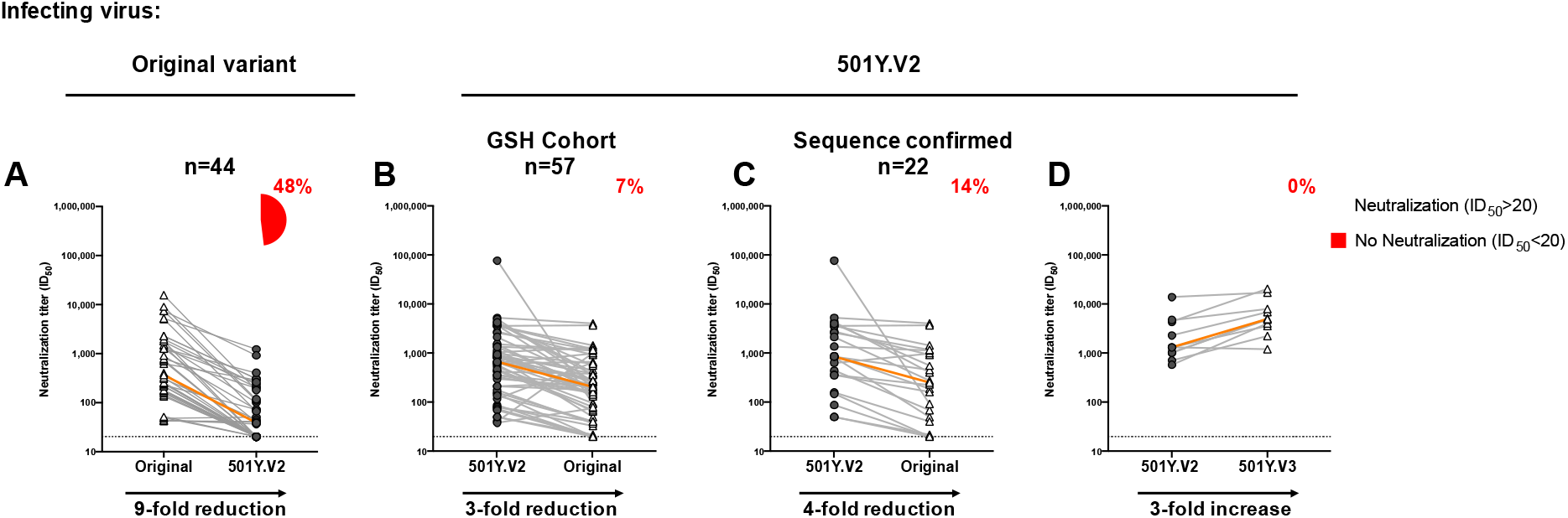
Neutralizing antibodies elicited by 501Y.V2 infection are more cross-reactive than those from patients infected with the original variant. (**A**) Plasma samples from patients infected with the original variant and (**B**-**C**) 501Y.V2-infected GSH cohort samples were compared for their neutralization cross-reactivity against other variants (n=57). In (**C**), the analysis was limited to those samples where sequencing confirmed infection by 501Y.V2 (n=22). (**D**) A subset of samples (n=10) was assayed against 501Y.V3 pseudoviruses. The orange line indicates the slope between the median neutralization potency of the samples tested. In the pie charts, purple indicates the proportion of samples with neutralization activity and red the proportion of samples with no detectable neutralization activity. The threshold of detection for the neutralization assay is ID_50_>20. All experiments were performed in duplicate. Data for the original variant plasma was taken from *Wibmer et al*., *2021, Nature Medicine*.

We also compared titers from wave 1, where infection was by the original variant (May to September 2020) and the wave 2 GSH plasma (501Y.V2 infection, December 2020 to January 2021) against both the original variant (**Figure S3A**) and 501Y.V2 (**Figure S3B**). The original variant was neutralized by both wave 1 and wave 2 sera (GMT: 510 and 203, respectively). However, the 501Y.V2 variant was neutralized 11-fold better by the wave 2 sera (GMT: 618) than wave 1 sera (GMT: 54) which was obtained from both hospitalized and non-hospitalized individuals **(Figure S3A, B)**.

## Discussion

Neutralizing antibodies are widely assumed to be a correlate of protection from SARS-CoV-2 infection (Folegatti et al., 2020, Jackson et al., 2020, Rogers et al., 2020). However, the emergence of multiple SARS-CoV-2 variants exhibiting similar escape substitutions suggests convergent evolution towards a more resistant phenotype, with major implications for the efficacy of vaccines globally (Fontanet et al., 2021). We and others have shown that neutralizing antibodies elicited by the original D614G variant failed to effectively neutralize 501Y.V2 (Wibmer et al., 2021, Cele et al., 2021). Here we show that, in contrast, antibodies elicited by 501Y.V2 neutralize both the original variant and 501Y.V3 (P.1), indicating high levels of cross-reactivity. These data suggest that vaccines based on the 501Y.V2 genetic sequence may be more broadly effective.

The level of neutralizing antibodies required for protection from SARS-CoV-2 infection has yet to be established. We show that titers of neutralizing antibodies elicited by the 501Y.V2 variant are relatively high (GMT: 618). It should be noted, however, that we used hospitalized participants who tend to mount higher neutralizing responses than individuals with mild disease (Seow et al., 2020). Since the majority of infections are mild/moderate, understanding both the cross-reactivity and durability of lower titer responses in 501Y.V2 infection will be important for predicting re-infection risk. Furthermore, studies of the cross-reactivity of the immune responses raised to other variants remain to be performed. Plasma from individuals infected with the 501Y.V1 variant (B.1.1.7), first detected in the UK, show reduced titers against 501Y.V2 suggesting this spike does not elicit particularly cross-reactive antibodies (Faulkner et al., 2021). This is similar to observations using samples from individuals infected with the original variant that lost neutralizing activity against 501Y.V2 (Wibmer et al., 2021, Cele et al., 2021). Cross-reactivity may be specific to variants with sequence or structural features similar to 501Y.V2, and therefore, studies of the immunity to other variants such as 501Y.V3 (P.1) will provide important additional information.

The targets of cross-reactive neutralizing antibodies elicited by 501Y.V2 remain to be defined. Stamatatos and colleagues showed that cross-reactive antibodies primed by SARS-CoV-2 infection and boosted with an mRNA vaccine neutralized both the original variant and 501Y.V2, through antibodies to the RBD and S2 but not the NTD (Stamatatos et al., 2021). Our data also suggests that much of the neutralizing response to 501Y.V2 is mediated by RBD-directed antibodies given that the neutralization profiles of the complete 501Y.V2 and the 501Y.V2 RBD triple mutant are similar. Although 501Y.V2 is resistant to class I/II RBD-directed antibodies, class III and IV RBD-specific antibodies are not affected by the RBD changes in the 501Y.V2 variant (Yuan et al., 2021, Wang et al., 2021a, Wang et al., 2021b). These classes represent a minority of RBD-specific antibodies against the original variant but may be more common in 501Y.V2 infections (Gavor et al., 2020, Liu et al., 2020, Wang et al., 2021b, Wibmer et al., 2021). Furthermore, an optimized SARS-CoV mAb which targets a highly conserved region on SARS-CoV-2 that overlaps with the ACE-2 binding site identifies another site targeted by cross-reactive antibodies (Rappazzo et al., 2021). The epitope of this mAb does not contact residues E484 and K417 but lies between the epitopes of Class I and Class IV RBD-specific antibodies (Rappazzo et al., 2021). Such specificities, if elicited during infection, may be the target of the antibody response elicited by the 501Y.V2 variant. Lastly, neutralizing antibodies recognizing the S2 subunit have been identified (Huang et al., 2021) and though these are less common in infection by the original variant than RBD-and NTD-specific antibodies, this site may also be a target for antibodies induced by 501Y.V2 infection.

Binding antibodies elicited by 501Y.V2 also show good cross-reactivity against original variant, a similar finding to our study on antibodies elicited by the original variant (Wibmer et al., 2021). This highlights the polyclonal nature of the binding antibody response to SARS-CoV-2 infection. A number of studies have demonstrated that effector functions of binding antibodies elicited during infection and vaccination contribute to protection from re-infection and severe disease (Yu et al., 2020, Atyeo et al., 2020). The role of these non-neutralizing antibodies and the efficacy of T cell responses elicited by the 501Y.V2 variant are unknown, but the significant proportion of cross-reactive antibodies elicited by both the original variant and 501Y.V2 is promising for vaccine design and protection from re-infection.

One limitation of our study is the absence of matched SARS-CoV-2 spike sequence data for all participants. However, of the 28 samples that were sequenced, all were 501Y.V2. The fact that similar findings were seen in the subset of those who were sequenced, also supports our conclusions (Tegally et al., 2021). This together with the finding that 501Y.V2 accounted almost all of the infections in South Africa, confirms the high probability that most infections in this cohort were caused by 501Y.V2. Furthermore, we have ruled out prior symptomatic infection by the original variant in all but 2 individuals. Although we cannot rule out prior asymptomatic infection, the high number of samples tested makes the probability of a substantial proportion of these being re-infections unlikely.

Overall, the emergence of variants with increased resistance to neutralizing antibodies has major implications for vaccine design. However, 501Y.V2 appears to generate a robust antibody response in hospitalized individuals. Our data indicate that vaccines built upon the spike protein of 501Y.V2 may be promising candidates for the elicitation of cross-reactive neutralizing antibodies to SARS-CoV-2.

## Materials and Methods

### Cohort

Plasma samples were obtained from hospitalized COVID-19 patients (n=89) with moderate disease admitted to Groote Schuur Hospital (GSH) cohort, Cape Town from 30 December 2020 – 15 January 2021 during the second wave in South Africa. All patients were aged ≥ 18 years and were HIV negative. This study received ethics approval from the Human Research Ethics Committee of the Faculty of Health Sciences, University of Cape Town (R021/2020). All patients had polymerase chain reaction (PCR) confirmed SARS-CoV-2 infection a median of 7 days (IQR 3-11) prior to blood collection. Clinical folders were consulted for 87/89 participants and none showed evidence of prior symptomatic COVID-19 disease.

We also used data from a previously described cohort infected with the original D614G variant during the first wave, for comparison (Wibmer et al., 2021). The Pretoria COVID study cohort consisted of adults (age > 18 years) who had PCR-confirmed SARS-CoV-2 infections, with moderate to severe COVID-19, and admitted to Steve Biko Academic Hospital (Pretoria, South Africa) from April to September 2020. Blood samples were collected at admission and seven days later, or at discharge from hospital (whichever was sooner). Ethics approval was received from the University of Pretoria, Human Research Ethics Committee (Medical) (247/2020).

### SARS-CoV-2 spike genome sequencing

RNA sequencing was performed as previously published (Pillay et al., 2020). Briefly, RNA extracted from swabs obtained from 28 GSH cohort participants was used to synthesize cDNA using the Superscript IV First Strand synthesis system (Life Technologies, Carlsbad, CA) and random hexamer primers. SARS-CoV-2 whole genome amplification was performed by multiplex PCR using primers designed on Primal Scheme (http://primal.zibraproject.org/) to generate 400 bp amplicons with a 70 bp overlap covering the SARS-CoV-2 genome. PCR products were cleaned up using AMPure XP magnetic beads (Beckman Coulter, CA) and quantified using the Qubit dsDNA High Sensitivity assay on the Qubit 3.0 instrument (Life Technologies Carlsbad, CA). The Illumina^®^ DNA Prep kit was used to prepare indexed paired end libraries of genomic DNA. Sequencing libraries were normalized to 4 nM, pooled and denatured with 0.2 N sodium hydroxide. Libraries were sequenced on the Illumina MiSeq instrument (Illumina, San Diego, CA, USA). Paired-end fastq reads were assembled using Genome Detective 1.132 (https://www.genomedetective.com) and the Coronavirus Typing Tool (Cleemput et al., 2020). The initial assembly obtained from Genome Detective was polished by aligning mapped reads to the references and filtering out low-quality mutations using bcftools 1.7-2 mpileup method. Mutations were confirmed visually with bam files using Geneious software (Biomatters Ltd, New Zealand).

### Clade and Lineage classification and phylogenetic analysis

To assign the sequenced samples to their lineage and clade, we used the dynamic lineage classification method proposed by Rambault et al. via the Phylogenetic Assignment of named Global Outbreak LINeages (PANGOLIN) software suite (https://github.com/hCoV-2019/pangolin) (Rambaut et al., 2020a), and Nextclade (Hadfield et al., 2018), respectively. We confirmed clade classification by analyzing our sequenced genomes against a global reference dataset using a custom pipeline based on a local version of NextStrain (https://github.com/nextstrain/ncov) (Hadfield et al., 2018). The pipeline contains several python scripts that manage the analysis workflow. It performs alignment of genomes in MAFFT (Katoh and Standley, 2013), phylogenetic tree inference in IQ-Tree V1.6.9 (Nguyen et al., 2015), tree dating and ancestral state construction and annotation (https://github.com/nextstrain/ncov). The phylogenetic trees were visualized using ggplot and ggtree (Yu et al., 2017, Wickham, 2016).

### Expression and purification of SARS-CoV-2 antigens

SARS-CoV-2 spike and SBD (RBD + subdomain 1) proteins were expressed in Human Embryonic Kidney (HEK) 293F suspension cells by transfecting the cells with SARS CoV-2 plasmid DNA. After incubating for six days at 37 °C, 70% humidity and 10% CO_2_, proteins were first purified using a nickel resin followed by size-exclusion chromatography. Relevant fractions were collected and frozen at -80 °C until use.

### Enzyme-linked Immunosorbent Assay (ELISA)

Two μg/ml of spike or SBD proteins were used to coat 96-well, high-binding plates and incubated overnight at 4 °C. The plates were blocked in blocking buffer consisting of 5% skimmed milk powder, 0.05% Tween 20, 1X PBS. Plasma samples were diluted to 1:100 starting dilution in blocking buffer and added to the plates. Secondary antibody was diluted to 1:3000 in blocking buffer and added to the plates followed by TMB substrate (Thermofisher Scientific). Upon stopping the reaction with 1M H_2_SO_4_, absorbance was measured at a 450nm wavelength. In all instances, mAbs CR3022 or palivizumab were used as positive and negative controls, respectively.

### SARS-CoV-2 pseudovirus based neutralization assay

SARS-CoV-2 pseudotyped lentiviruses were prepared by co-transfecting the HEK 293T cell line with either the SARS-CoV-2 614G spike (D614G), SARS-CoV2 501Y.V2-RBD spike (K417N, E484K and N501Y, D614G) or SARS-CoV-2 501Y.V2 spike (L18F, D80A, D215G, K417N, E484K, N501Y, D614G, A701V, 242-244 del) and the SARS-CoV-2 501Y.V3 spike (L18F, T20N, P26S, D138Y, R190S, K417T, E484K, N501Y, D614G, H655Y, T1027I, V1176F) plasmids in conjunction with a firefly luciferase encoding lentivirus backbone plasmid and a murine leukemia virus backbone plasmid. The parental plasmids were kindly provided by Drs Elise Landais and Devin Sok (IAVI). For the neutralization assay, heat-inactivated plasma samples from COVID-19 convalescent donors were incubated with the SARS-CoV-2 pseudotyped virus for 1 hour at 37°C, 5% CO_2_. Subsequently, 1×10^4^ HEK293T cells engineered to over-express ACE-2, kindly provided by Dr Michael Farzan (Scripps Research), were added and incubated at 37°C, 5% C0_2_ for 72 hours upon which the luminescence of the luciferase gene was measured. CB6, CA1, CR3022 and palivizumab were used as controls.

## Acknowledgements

We thank the participants in the GSH cohort and staff who helped in the collection of samples. Mieke van der Mescht, Zelda van der Walt, Talita de Villiers, Fareed Abdullah, Paul Rheeder, Albertus Malan, Simon Spoor and Wesley van Hougenhouck-Tulleken are thanked for contributing to patient management, sample collection and processing, and data management for the Pretoria COVID study. We thank Richard Baguma, Roanne Keeton and Marius Tincho for sample processing and shipping. We thank Stephen Korsman and Tamryn Smith for assistance with sample identification and access. We acknowledge funding from the South African Medical Research Council (Reference numbers 96825, SHIPNCD 76756 and DST/CON 0250/2012), United States Centers for Disease Control and Prevention (Grant number 5 U01IP001048-05-00), the ELMA South Africa Foundation (Grant number 20-ESA011) and the Wellcome Centre for Infectious Diseases Research in Africa (CIDRI-Africa), which is supported by core funding from the Wellcome Trust [203135/Z/16/Z]. P.L.M. is supported by the South African Research Chairs Initiative of the Department of Science and Innovation and the NRF (Grant No 9834). C.K.W. is supported by Fogarty International Center of the National Institutes of Health under Award Number R21TW011454 as well as the FLAIR Fellowship program under award number FLR\R1\201782. KvdB is supported in part by the Fogarty International Centre or the National Institutes of Health under Award Number 1D43TW010345. W.A.B. is supported by the EDCTP2 programme of the European Union (EU)’s Horizon 2020 programme (TMA2016SF-1535-CaTCH-22). We thank Drs Nicole Doria-Rose, David Montefiori, Elise Landais and Michael Farzan for reagents and assistance in establishing the SARS-CoV-2 pseudotyped neutralization assay and enabling equivalency and proficiency testing. We thank Drs Devin Sok, Elise Landais, Dennis Burton, Nicole Doria-Rose and Peter Kwong for SARS-CoV-2 directed mAbs. We are grateful to Jason McLellan for the WT Hexapro Spike construct. We thank Anne von Gottberg, Cheryl Cohen and Susan Meiring for samples in the previously published comparator group. We thank the informal 501Y.V2 consortium of South African scientists, chaired by Drs Willem Hanekom and Tulio de Oliveira for suggestions and discussion of data, and Drs Valerie Mizrahi and Robert Wilkinson for enabling this collaboration. We also thank all NGS-SA laboratories in South Africa that were responsible for producing the SARS-CoV-2 genomes that enabled the rapid dissemination of SARS-CoV-2 sequences.

