## Supplementary Data for "SARS-CoV-2 501Y.V2 (B.1.351) elicits cross-reactive neutralizing antibodies"

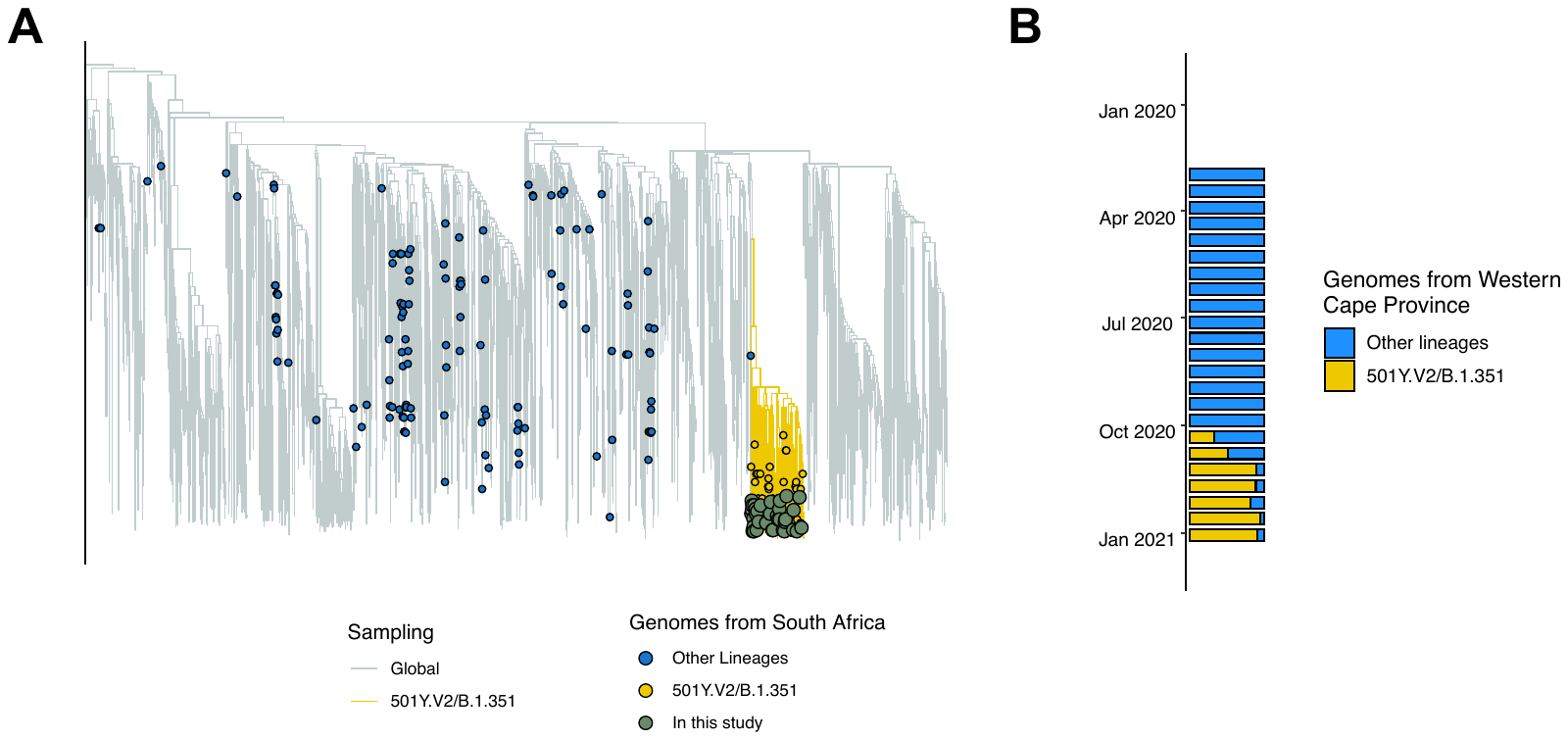

**Figure S1. SARS-CoV-2 genome sequencing confirms dominance of infection by the 501Y.V2 variant in Western Cape, South Africa.** Genomic sequencing was conducted in the Western Cape Province of South Africa (the same province where our GSH cohort is based) on an approximately weekly basis from the second week of March 2020 to the second week of January 2020. Swabs from 28 individuals from the GSH cohort were sequenced to confirm the infecting variant. (**A**) A time-resolved maximum clade credibility phylogeny of 2621 SARS-CoV-2 sequences, 209 of which are from South Africa and denoted with tip points. The 501Y.V2 (B.1.351) cluster is highlighted in yellow, and the genomes from this study are shown in green, all falling in that cluster. One genome, although assigned to the 501Y.V2/B.1.351 lineage by mutations, was excluded from the phylogenetic tree because of low coverage. (**B**) The bar plot shows the progressive proportion of the 501Y.V2/B.1.351 lineage in Western Cape.

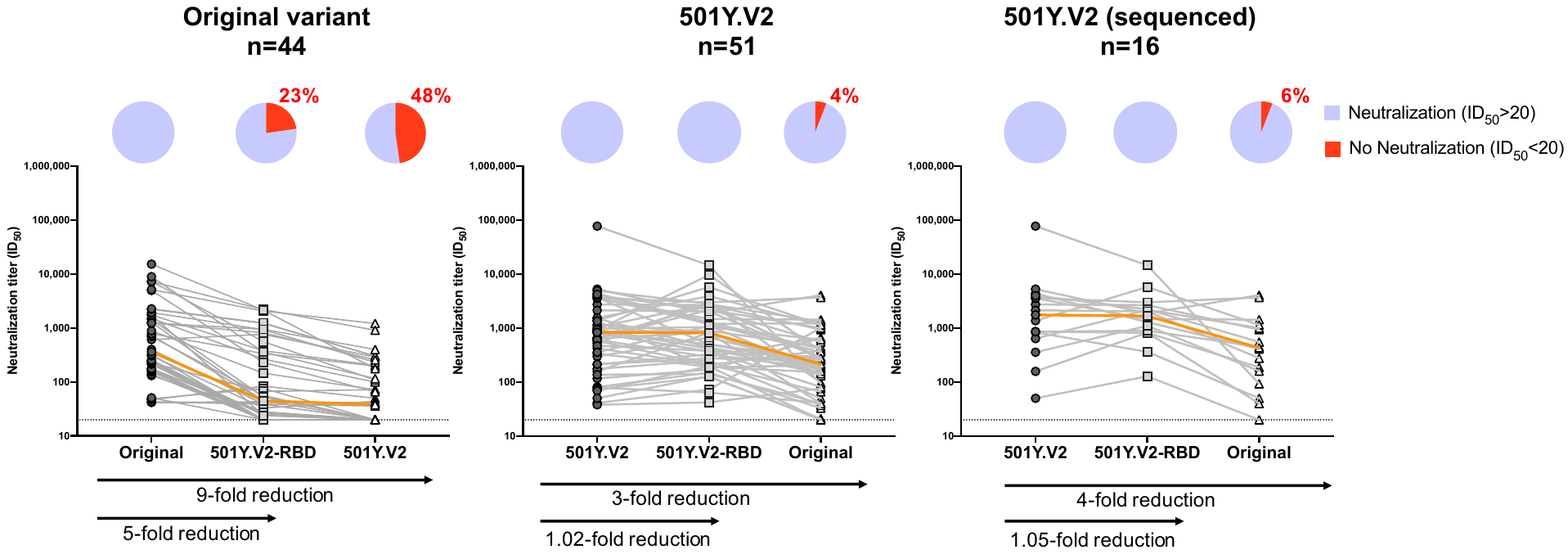

**Figure S2.** **Antibody cross-reactivity against the full 501Y.V2, 501Y.V2-RBD and the original variant in plasma from patients infected with 501Y.V2.** (**A**) Plasma samples from patients infected with the original variant and (**B**-**C**) 501Y.V2-infected GSH cohort samples were compared for their neutralization cross-reactivity against 501Y.V2, 501Y.V2-RBD and D614G (n=51). In (**C**), the analysis was limited to those samples where sequencing confirmed infection by 501Y.V2 (n=16). The orange line indicates the slope between the median neutralization potency of the samples tested. In the pie charts, purple indicates the proportion of samples with neutralization activity and red the proportion of samples with no detectable neutralization activity. The threshold of detection for the neutralization assay is ID_50_>20. All experiments were performed in duplicate. Data for the original virus plasma was taken from *Wibmer et al., 2021, Nature Medicine*.

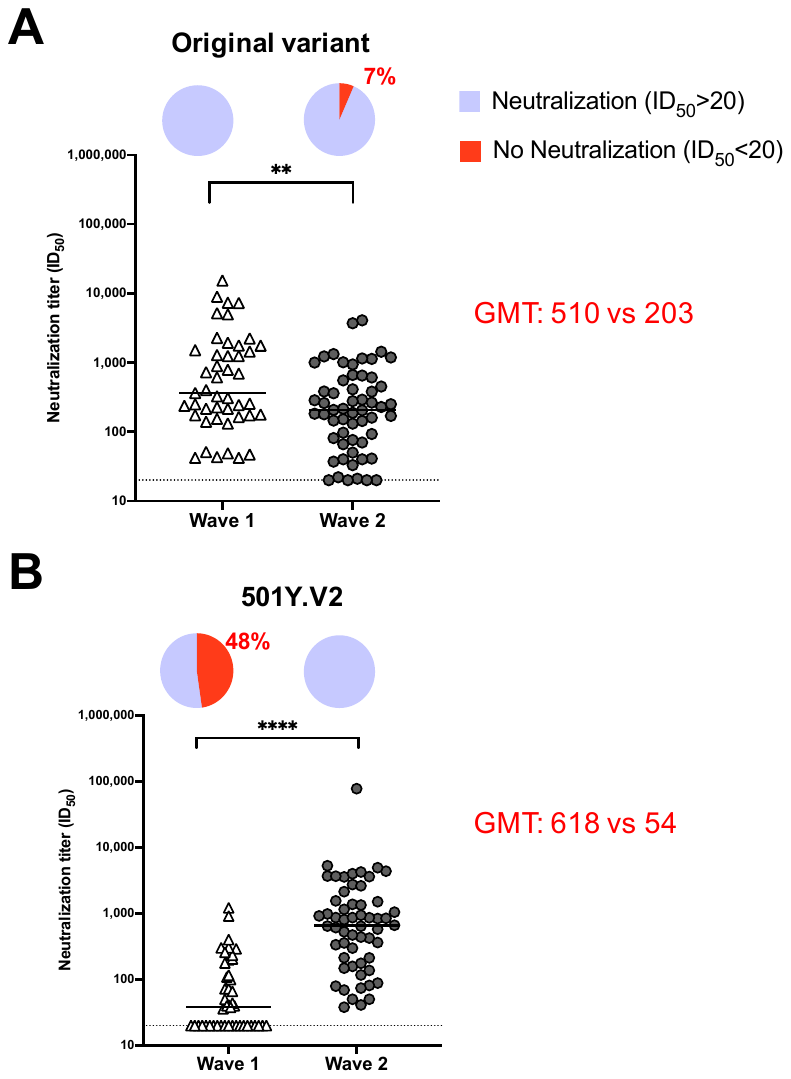

**Figure S3. Plasma elicited by 501Y.V2 infection (Wave 2) is more cross-reactive than plasma from the original virus infection (Wave 1).** Plasma from patients infected with 501Y.V2 or the original variant were tested against (**A**) the original virus or (**B**) the 501Y.V2 variant. In the pie charts, purple indicates the proportion of samples with neutralization activity and red the proportion of samples with no detectable neutralization activity. The threshold of detection for the neutralization assay is ID_50_>20. All experiments were performed in duplicate. Data for the original virus plasma was taken from *Wibmer et al., 2021, Nature Medicine*. Significance is shown as: **p<0.01 and ****p<0.0001.
